# Dopamine modulates prediction error forwarding in the nonlemniscal inferior colliculus

**DOI:** 10.1101/824656

**Authors:** Catalina Valdés-Baizabal, Guillermo V. Carbajal, David Pérez-González, Manuel S. Malmierca

## Abstract

The predictive processing framework describes perception as a hierarchical predictive model of sensation. Higher-level neural structures constrain the processing at lower-level structures by suppressing synaptic activity induced by predictable sensory input. But when predictions fail, deviant input is forwarded bottom-up as ‘prediction error’ to update the perceptual model. The earliest prediction error signals identified in the auditory pathway emerge from the nonlemniscal inferior colliculus (IC). The drive that these feedback signals exert on the perceptual model depends on their ‘expected precision’, which determines the postsynaptic gain applied in prediction error forwarding. Expected precision is theoretically encoded by the neuromodulatory (e.g., dopaminergic) systems. To test this empirically, we recorded extracellular responses from the rat nonlemniscal IC to oddball and cascade sequences before, during and after the microiontophoretic application of dopamine or eticlopride (a D2-like receptor antagonist). Hence, we studied dopaminergic modulation on the subcortical processing of unpredictable and predictable auditory changes. Results demonstrate that dopamine reduces the net neuronal responsiveness exclusively to unexpected input, without significantly altering the processing of expected auditory events at population level. We propose that, in natural conditions, dopaminergic projections from the thalamic subparafascicular nucleus to the nonlemniscal IC could serve as a precision-weighting mechanism mediated by D2-like receptors. Thereby, the levels of dopamine release in the nonlemniscal IC could modulate the early bottom-up flow of prediction error signals in the auditory system by encoding their expected precision.

## Introduction

Perceptual systems prune redundant sensory input as a means of sparing processing resources while providing saliency to input that is rare, unique, and therefore potentially more informative. This perceptual function has been classically studied in humans using the auditory oddball paradigm [1], where the successive repetition of a tone (‘standard condition’, henceforth ‘STD’) is randomly interrupted by an ‘oddball’ tone (‘deviant condition’, henceforth ‘DEV’). When applied to animal models, the oddball paradigm unveils a phenomenon of neuronal short-term plasticity called stimulus-specific adaptation (SSA), measured as the difference between DEV and STD responses [2].

SSA first emerges in the auditory system at the level of the inferior colliculus (IC), mainly in its nonlemniscal portion (i.e., the IC cortices) [3]. As a site of convergence of both ascending and descending auditory pathways, the IC plays a key role in processing deviant sounds over redundant ones [4] and shaping the auditory context [5]. The complex computational network of the IC integrates excitatory, inhibitory and rich neuromodulatory input [6,7]. This includes dopaminergic innervation from the subparafascicular nucleus (SPF) of the thalamus to the nonlemniscal IC [8–12]. Previous reports have detected mRNA coding for dopaminergic D2-like receptors in the IC [9,13] and proved its functional expression as protein [14], while other studies demonstrated that dopamine modulates the auditory responses of IC neurons in heterogeneous manners [10,14]. However, the involvement of dopaminergic modulation of SSA in the IC is yet to be confirmed.

In recent years, subcortical SSA in the auditory system has been reinterpreted in the context of the predictive processing [15]. This conceptual framework posits that hierarchically coupled neuronal populations infer the hidden causes of sensation and predict upcoming sensory regularities using a generative model of the world [16–25]. In such hierarchical model, higher neural populations try to explain away or inhibit the sensory input prompted by the hidden states of the world. As a result, lower-level neural populations receiving those top-down predictions decrease their responsiveness to expected sensory inputs, which during an oddball paradigm manifest functionally as SSA of the STD response. But when encountering a DEV, the generative model fails to predict that ‘oddball’, forwarding a prediction error (PE) signal which reports the unexpected portions of sensory input to the higher-level neural population. That bottom-up flow of PE signals serves to provide feedback and update the inferred representations about the states of the world along each level of the processing hierarchy. In a previous study from our lab performed in awake and anaesthetized rodents, we demonstrated that DEV responses in the nonlemniscal IC were better explained as PE signaling activity [26].

In the predictive processing framework, there are only two sorts of things that need to be inferred about the world: the state of the world, and the uncertainty about that state [27]. On the one hand, representations about the states of the world emerge from the hierarchical exchange of top-down predictions and bottom-up PEs, which is embodied in the synaptic activity of the nervous system. On the other hand, this inferential process entails a certain degree of uncertainty, which is encoded in terms of *expected precision* or confidence by means of the postsynaptic gain [27–29]. Thereby, synaptic messages are weighted according to their expected precision as they are passed along the processing hierarchy. When expected precision is high, PE signals receive postsynaptic amplification to strengthen its updating power. Conversely, when imprecision is expected, PE signals undergo negative gain to prevent misrepresentations. Neuromodulators (such as dopamine) cannot directly excite or inhibit postsynaptic responses, but only modulate the postsynaptic responses to other neurotransmitters. Therefore, according to some predictive processing implementations [27,30–32], the only possible function of neuromodulatory systems is to encode the expected precision.

In this study, we perform microiontophoretic injections of dopamine and eticlopride (a D2-like receptor antagonist) while recording single- and multi-unit responses under oddball and regular sequences to determine whether dopaminergic input to the nonlemniscal IC modulates response properties and predictive processing. Our results demonstrate that dopamine has a profound effect on how unexpected sounds are processed in the nonlemniscal IC. We argue this could be compatible with a dopaminergic encoding of the expected precision of PE signals, acting as a regulatory mechanism at the level of the auditory brainstem.

## Materials and Methods

### Surgical procedures

We conducted experiments on 31 female Long-Evans rats aged 9–17 weeks with body weights between 150–250 gr. All methodological procedures were approved by the Bioethics Committee for Animal Care of the University of Salamanca (USAL-ID-195), and performed in compliance with the standards of the European Convention ETS 123, the European Union Directive 2010/63/EU and the Spanish Royal Decree 53/2013 for the use of animals in scientific research.

We first induced surgical anesthesia with a mixture of ketamine/xylazine (100 and 20 mg/Kg respectively, intramuscular) and then maintained it with urethane (1.9 g/Kg, intraperitoneal). To ensure a stable deep anesthetic level, we administered supplementary doses of urethane (~0.5 g/Kg, intraperitoneal) when the corneal or pedal withdrawal reflexes were present. We selected urethane over other anesthetic agents because it better preserves normal neural activity, having a modest, balanced effect on inhibitory and excitatory synapses [33–36].

Prior to the surgery, we recorded auditory brainstem responses (ABR) with subcutaneous needle electrodes to verify the normal hearing of the rat. We acquired the ABR using a RZ6 Multi I/O Processor (Tucker-Davis Technologies, TDT) and BioSig software (TDT) before beginning each experiment. ABR stimuli consisted of 0.1 ms clicks at a rate of 21 clicks/s, delivered monaurally to the right ear in 10 dB steps, from 10 to 90 decibels of sound pressure level (dB SPL), in a closed system through a speaker coupled to a small tube sealed in the ear.

After normal hearing was confirmed, we placed the rat in a stereotaxic frame where the ear bars were replaced by hollow specula that accommodated the sound delivery system. We performed a craniotomy in the left parietal bone to expose the cerebral cortex overlying the left IC. We removed the dura overlying the left IC and covered the exposed cortex with 2% agar to prevent desiccation.

### Data acquisition procedures

Experiments were performed inside a sound-insulated and electrically shielded chamber. All sound stimuli were generated using a RZ6 Multi I/O Processor (TDT) and custom software programmed with OpenEx Suite (TDT) and MATLAB (MathWorks). In search of evoked auditory neuronal responses from the IC, we presented white noise bursts and sinusoidal pure tones of 75 ms duration with 5 ms rise-fall ramps. Once the activity of an auditory unit was clearly identified, we only used pure tones to record the experimental stimulation protocols. All protocols ran at 4 stimuli per second and were delivered monaurally in a closed-field condition to the ear contralateral to the left IC through a speaker. We calibrated the speaker using a ¼-inch condenser microphone (model 4136, Brüel&Kjær) and a dynamic signal analyzer (Photon+, Brüel&Kjær) to ensure a flat spectrum up to ~73 dB SPL between 0.5 and 44 kHz, and that the second and third signal harmonics were at least 40 dB lower than the fundamental at the loudest output level.

To record extracellular activity while carrying out microiontophoretic injections, we attached a 5-barrel glass pipette to a hand-manufactured, glass-coated tungsten microelectrode (1.4–3.5 MΩ impedance at 1 kHz), with the tip of the electrode protruding 15–25 μm from the pipette tip [37]. We place the electrode over the exposed cortex, forming an angle of 20° with the horizontal plane towards the rostral direction. Using a piezoelectric micromanipulator (Sensapex), we advanced the electrode while measuring the penetration depth until we could observe a strong spiking activity synchronized with the train of searching stimuli.

Analog signals were digitized with a RZ6 Multi I/O Processor, a RA16PA Medusa Preamplifier and a ZC16 headstage (TDT) at 12 kHz sampling rate and amplified 251×. Neurophysiological signals for multiunit activity were band-pass filtered between 0.5 and 4.5 kHz. Stimulus generation and neuronal response processing and visualization were controlled online with custom software created with the OpenEx suite (TDT) and MATLAB. A unilateral threshold for automatic action potential detection was manually set at about 2-3 standard deviations of the background noise. Spike waveforms were displayed on the screen and overlapped on each other in a pile-plot to facilitate isolation of units. Recorded spikes were considered to belong to a single unit when the SNR of the average waveform was larger than 5 (51% of the recorded units).

### Stimulation protocols

For all recorded neurons, we first computed the frequency-response area (FRA), which is the map of response magnitude for each frequency/intensity combination (Fig. 1). The stimulation protocol to obtain the FRA consisted of a randomized sequence of sinusoidal pure tones ranging between 0.7-44 kHz, 75 ms of duration with 5 ms rise-fall ramps, presented at a 4 Hz rate, randomly varying frequency and intensity of the presented tones (3–5 repetitions of all tones).

**Figure 1.**
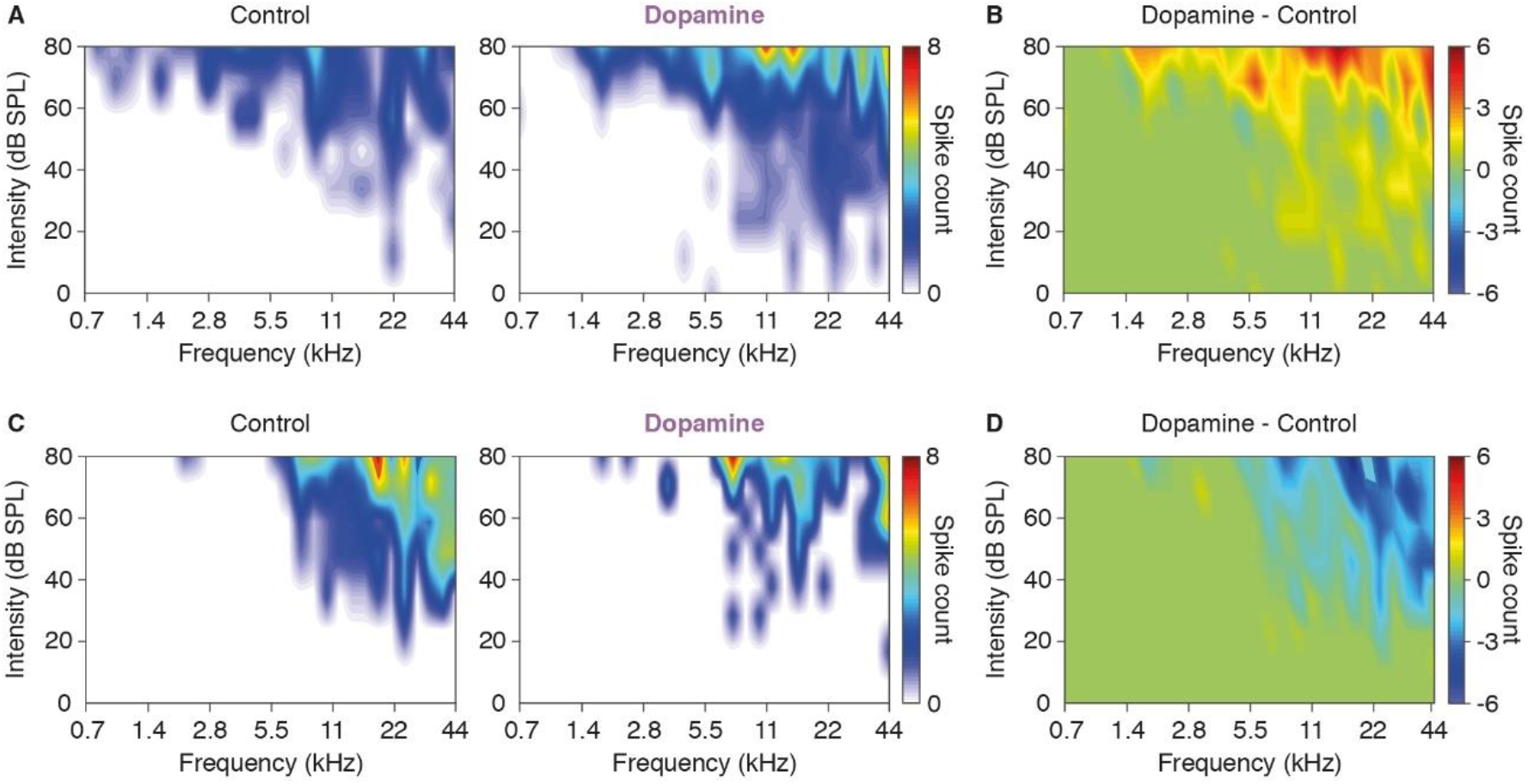
Effect of dopamine on the FRA. **A)** FRA of a neuron in control condition (left panel) and after dopamine application (right panel). **B)** The subtraction of the control FRA from that during application of dopamine in A reveals that dopamine increased the excitability of this neuron. **C)** FRA of another neuron in control condition (left panel) and after dopamine application (right panel). **D)** The subtraction of the control FRA from that during application of dopamine in C reveals that dopamine decreased the excitability of this neuron.

### Protocol 1: Oddball paradigm (DEV and STD)

In a first round of experiments, we used the oddball paradigm (Fig. 4A) to study SSA. We presented trains of 400 stimuli containing two different frequencies (*f_1_* and *f_2_*) in a pseudo-random order at a 4Hz repetition rate and at a level of 10–40 dB above threshold. Both frequencies were within the excitatory FRA previously determined for the neuron (Fig. 1), and evoked similar firing rates (FR). One frequency (*f_1_*) appeared with high probability within the sequence (STD; P=0.9). The succession of STD tones was randomly interspersed with the second frequency (*f_2_*), presented with low probability within the sequence (DEV; P=0.1). After obtaining one data set, the relative probabilities of the two stimuli were reversed, with *f_2_* becoming the STD and *f_1_* becoming the DEV (Fig. 4A). This allows to control for the physical characteristics of the sound in the evoked response, such that the differential response between DEV and STD of a given tone can only be due to their differential probability of appearance. The separation between *f_1_* and *f_2_* was 0.28 (49 units) or 0.5 (45 units; Protocol 2 was also applied to these units) octaves, which is within the range of frequency separations used in other previous studies [3,26,38–43]. The units from those two groups were pooled together, since their responses did not differ significantly. Deviant and standard responses were averaged from all stimulus presentations from both tested frequencies.

The Common SSA Index (CSI) was calculated as:

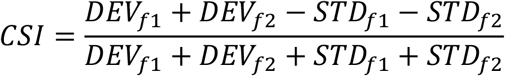

where *DEV_fi_* and *STD_fi_* are FRs in response to a frequency *f_i_* when it was presented in deviant and standard conditions, respectively. The CSI ranges between −1 to +1, being positive if the DEV response was greater than the STD response. The firing rates in response to DEV or STD stimuli were calculated using 100 ms windows starting at the beginning of each stimulus. Spontaneous firing rates were calculated using 75 ms windows previous to each individual stimulus.

### Protocol 2: Oddball paradigm + Cascade sequence (CAS)

In light of the recent discovery of PE signals recorded in the nonlemniscal IC [26], we decided to adapt our stimulation protocol to that of Parras and colleagues for a second round of experiments which incorporated the cascade sequence (CAS) [44]. By arranging a set of 10 tones in a regular succession of ascending or descending frequency, no tone is ever immediately repeated. Consequently, whereas CAS does not induce SSA—as opposed to STD—, its pattern remains predictable, so the next tone in the sequence can be expected— as opposed to DEV. Thus, this design contains 3 conditions of auditory transit: (1) *no change* or *predictable repetition* (i.e., STD), which is the most susceptible to SSA or repetition suppression (Fig. 4A, bottom); (2) *predictable change* (i.e., CAS; Fig. 4B); and (3) *unpredictable change* (i.e., DEV), which should allegedly elicit the strongest PE signaling when it surprisingly interrupts the uniform train of STDs (Fig. 4A, top).

Therefore, after computing the FRA (Fig. 1), we selected 10 evenly-spaced tones at a fixed sound intensity 10–40 dB above minimal response threshold, so that at least two tones fell within the FRA limits. Those 10 frequencies were separated from each other by 0.5 octaves, in order to make the results comparable to those of [26]. We used the 10 tones to build the ascending and descending versions of CAS (Fig. 4B). We selected 2 tones within that lot to generate the ascending and descending versions of the oddball paradigm (Fig. 4A), comparing the resultant DEV with their corresponding CAS versions (Fig. 4B). All sequences were 400 tones in length, at the same, constant presentation rate of 4 Hz. Thus, each frequency could be compared with itself in DEV, STD and CAS conditions (Fig. 4A-B), obtaining 40 trials per condition. To allow comparison between responses from different neurons, we normalized the spike count evoked by each tone in DEV, STD and CAS as follows:

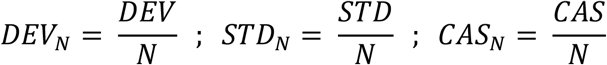

Where,

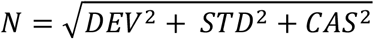

From these normalized responses, we computed the index of neuronal mismatch (iMM) as:

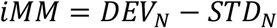

These indices range between −1 and 1. The iMM is largely equivalent to the classic CSI as an index of SSA, as previously demonstrated by Parras and colleagues (see their supplementary figure 2 in [26]). Nevertheless, please beware the CSI provides one index for each pair of tones in the oddball paradigm, whereas the iMM provides one index for each tone tested.

### Dopaminergic manipulation procedures

After recording the chosen stimulation protocol in a *‘control condition’*, i.e., before any dopaminergic manipulation, we applied either dopamine or the D2-like receptor antagonist eticlopride (Sigma-Aldrich Spain) iontophoretically through multi-barreled pipettes attached to the recording electrode. The glass pipette consisted of 5 barrels in an H configuration (World Precision Instruments, catalogue no. 5B120F-4) with the tip broken to a diameter of 30–40 μm [37]. The center barrel was filled with saline for current compensation (165 mM NaCl). The others were filled with dopamine (500 mM) or eticlopride (25 mM). Each drug was dissolved in distilled water and the acidity of the solution was adjusted with HCl to pH 3.5 for dopamine and pH 5 for eticlopride. The drugs were retained in the pipette with a −20 nA current and ejected using 90 nA currents (Neurophore BH-2 system, Harvard Apparatus). Thus, we released dopamine or eticlopride into the microdomain of the recorded neuron at concentrations that have been previously demonstrated effective in *in vivo* studies [10]. About 5 minutes after the drug injection, we repeated the FRA and the chosen stimulation protocol continuously until the drug was washed away, leaving roughly 2–3 minutes between the end of one recording set and the beginning of the next one. The recording set showing a maximal SSA alteration relative to the control values was considered the *‘drug condition’* of that neuron. We established the *‘recovery condition’* when the CSI returned to levels that did not significantly differ from control values, never before 40 minutes post-injection. We used either dopamine or eticlopride during protocol 1, while only dopamine was tested during protocol 2.

### Histological verification procedures

At the end of each experiment, we inflicted electrolytic lesions (5 μA, 5 s) through the recording electrode. Animals were sacrificed with a lethal dose of pentobarbital, after which they were decapitated, and the brains immediately immersed in a mixture of 1% paraformaldehyde and 1% glutaraldehyde in 1 M PBS. After fixation, tissue was cryoprotected in 30% sucrose and sectioned in the coronal plane at 40 μm thickness on a freezing microtome. We stained slices with 0.1% cresyl violet to facilitate identification of cytoarchitectural boundaries. Finally, we assigned the recorded units to one of the main subdivisions of the IC using the standard sections from a rat brain atlas as reference [45].

### Data analysis

The peristimulus histograms representing the time-course of the responses (Fig. 2E-F, 3E-F) were calculated using 1 ms bins an then smoothed with a 6 ms gaussian kernel (“ksdensity” function in MATLAB) to estimate the spike-density function over time.

**Figure 2.**
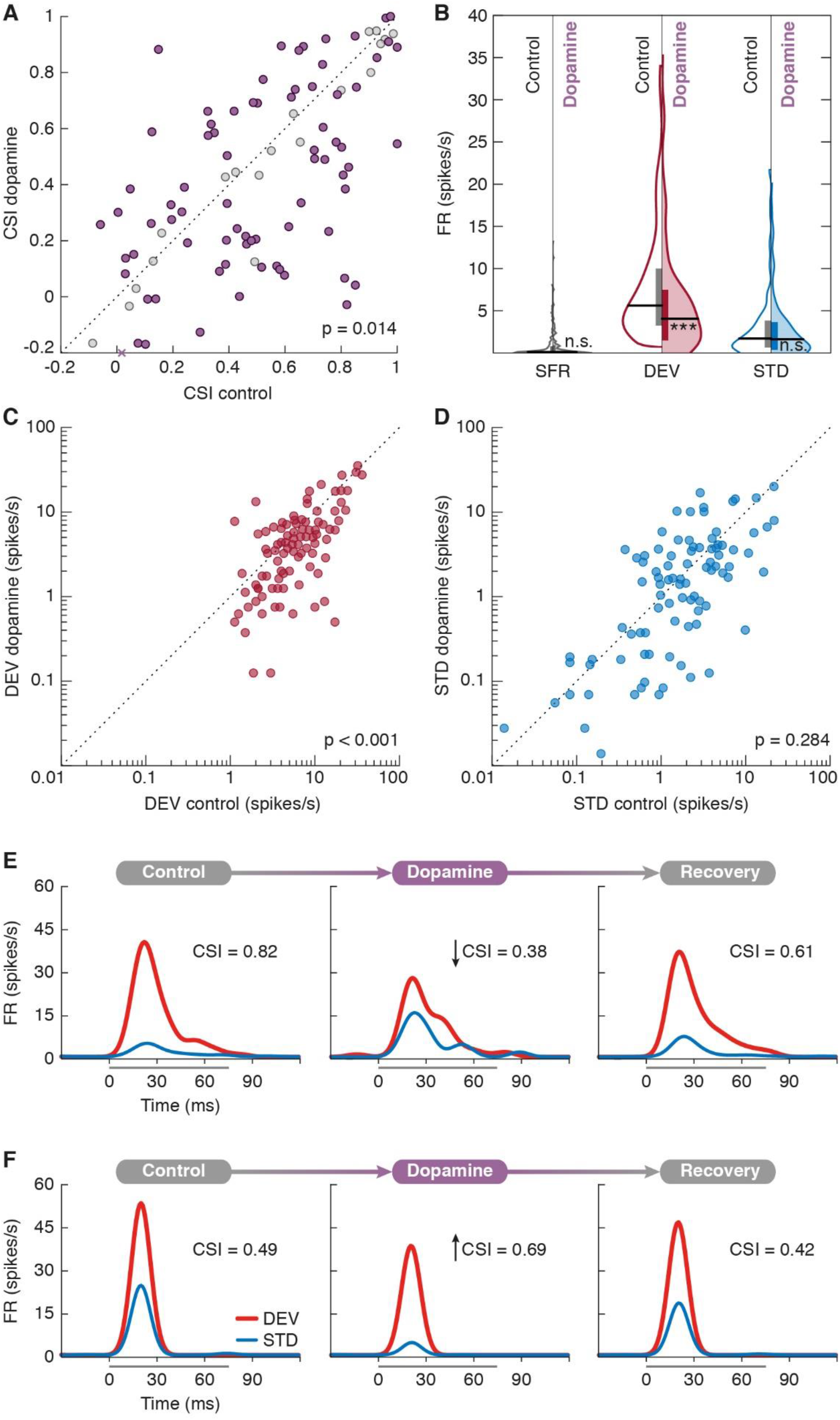
Effects of dopamine on the CSI. **A)** Scatter plot of the CSI in control condition versus dopamine application. Units that underwent significant CSI changes are represented in purple, whereas the rest are marked as gray dots. The purple cross on the abscissa axis represents one CSI measurement which ordinate value falls out of scale (y = −0.59). **B)** Violin plots of the SFR (gray), DEV response (red) and STD response (blue). Control conditions are represented in the left half of each violin (no color) while dopamine effects are on display in the right half (colored). Horizontal thick black lines mark the median of each distribution, while vertical bars cover the interquartile range. Regarding statistical significance, **n.s.** indicates that p > 0.05 and *** indicates that p < 0.001. **C)** Scatter plot of DEV responses in control condition versus dopamine application. **D)** Scatter plot of STD responses in control condition versus dopamine application. **E)** Peristimulus histogram of a unit before (left panel), during (middle panel) and after (right panel) dopamine application. In this case, dopamine reduced the CSI. **F)** Another example showing the opposite effects.

All the data analyses were performed with SigmaPlot (Systat Software) and MATLAB software, using the built-in functions, the Statistics and Machine Learning toolbox for MATLAB, as well as custom scripts and functions developed in our laboratory. Unless stated otherwise, all average values for trials and neurons in the present study are expressed as *‘median [interquartile range]’*, since the data did not follow a normal distribution (one-sample Kolmogorov-Smirnov test).

We performed a bootstrap procedure to analyze dopaminergic effects on each individual neuron. SSA indices, both CSI [2] and iMM [26], are calculated from the averages of the single-trial responses to DEV, STD and CAS. Consequently, only one value of such indexes can be obtained for each unit and condition. Therefore, to test the drug effects on each unit, we calculated the 95% bootstrap confidence intervals for the SSA index in the control condition. The bootstrap procedure draws random samples (with replacement) from the spike counts evoked on each trial, separately for DEV and STD stimuli, and then applies either the CSI or the iMM formula. This procedure is repeated 10000 times, thus obtaining a distribution of expected CSI values based on the actual responses from a single recording. We applied this procedure using the *bootci* MATLAB function, as in previous studies of SSA neuromodulation in the IC [38,39,42,46], which returned the 95% confidence interval for the CSI in the control condition. We considered drug effects to be significant when SSA index in the drug condition did not overlap with the confidence interval in the control condition.

We used the Wilcoxon Signed Rank test (*signrank* function in MATLAB) to check for differences at the population level between the control and drug CSI and FRs.

## Results

In order to study the role of dopamine in shaping SSA in the nonlemniscal IC, we recorded responses from a total of 142 single- and multi-units in 31 young adult Long Evans rats. In a first series of experiments (protocol 1, see *Methods*) we presented the oddball paradigm before, during and after microiontophoretic application of dopamine (n=94) or eticlopride (n=43). In an additional series of experiments (n=43, from the former pool of units), we also presented two cascade sequences (ascending and descending) in addition to the oddball paradigm [26] (protocol 2, see *Methods*). Histological verification located all recorded neurons in the rostral cortex of the IC, where SSA indexes tend to be higher [3,4,43].

### Dopamine effects on the CSI

The microiontophoretic application of dopamine caused an average 15% decrease of SSA in our sample, falling from a median CSI of 0.51 [0.25–0.78] in the control condition to 0.43 [0.15–0.73] after dopamine application (p=0.014; Figure 2A). Such average reduction was caused by a 26% drop in the median DEV response (control FR: 5.63 [3.25–10.00] spikes/s; dopamine FR: 4.19 [1.63–7.63]; p<0.001; red in Figure 2B,C), while the STD response did not show a significant change (control FR: 1.71 [0.64–3.83]; dopamine FR: 1.63 [0.36-3.65]; p=0.284; blue in Figure 2B,D). As observed in previous reports [3], the spontaneous firing rate (SFR) found in the nonlemniscal IC tended to be very scarce (for individual examples, see Figs. 2E-F and 3E-F) and did not change significantly with dopamine application (control SFR: 0.18 [0.05–0.77]; dopamine SFR: 0.15 [0.03–0.90]; p=0.525; gray in Figure 2B).

**Figure 3.**
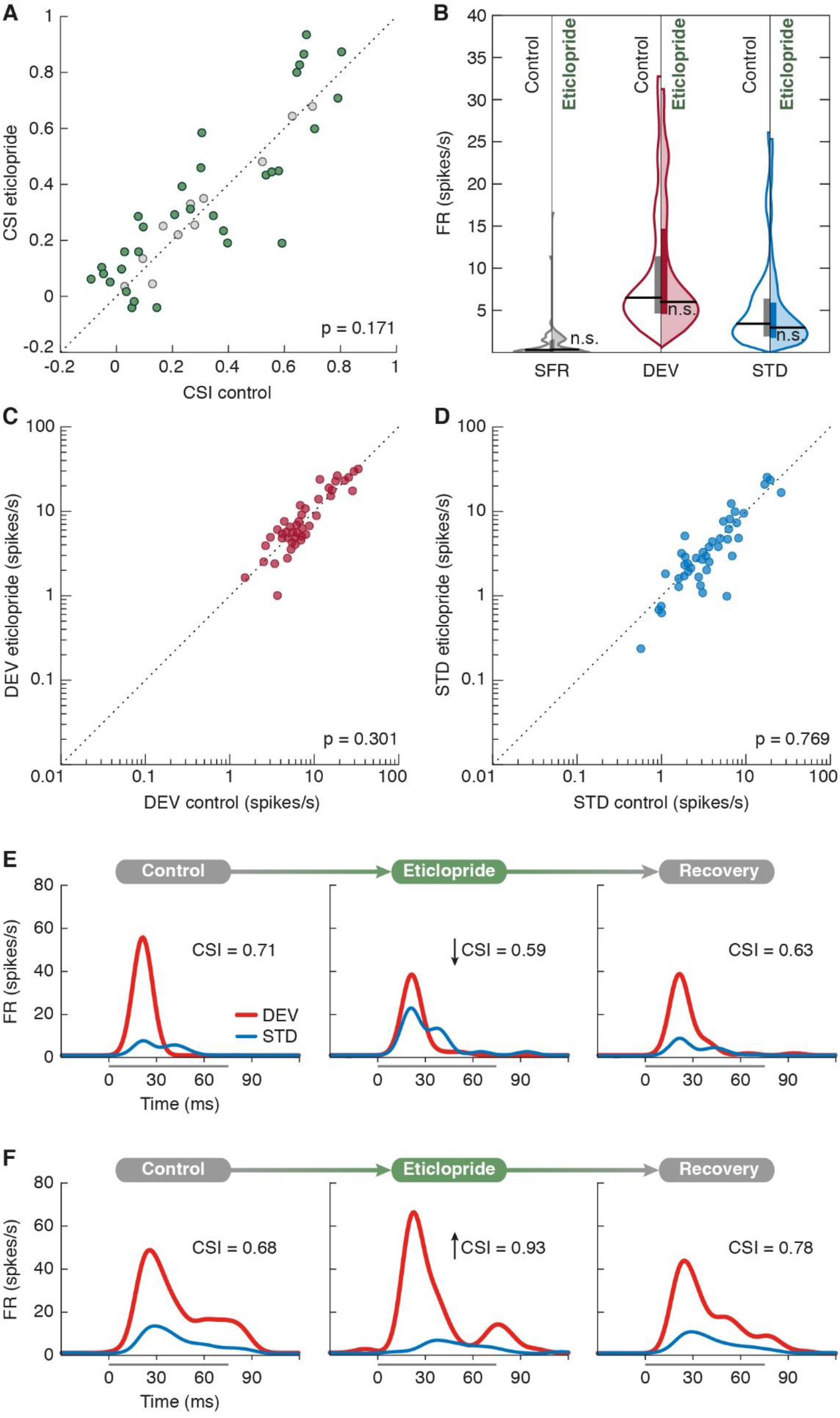
Effects of eticlopride on the CSI. **A)** Scatter plot of the CSI in control condition versus eticlopride application. Units that underwent significant CSI changes are represented in green, whereas the rest are marked as gray dots. **B)** Violin plots of the SFR (gray), DEV response (red) and STD response (blue). Control conditions are represented in the left half of each violin (no color) while eticlopride effects are on display in the right half (colored). Horizontal thick black lines mark the median of each distribution, while vertical bars cover the interquartile range. Regarding statistical significance, **n.s.** indicates that p > 0.05. **C)** Scatter plot of DEV responses in control condition versus eticlopride application. **D)** Scatter plot of STD responses in control condition versus eticlopride application. **E)** Peristimulus histogram of a unit before (left panel), during (middle panel) and after (right panel) eticlopride application. In this case, eticlopride reduced the CSI. **F)** Another unit example showing the opposite effects.

Previous studies had reported heterogeneous dopaminergic effects on the response of IC neurons [10,14], so we performed a bootstrap analysis to evaluate the statistical significance of CSI changes unit by unit. We confirmed such heterogeneity across our sample, with 42 units following the population trend by decreasing their CSI, whereas 33 units showed CSI increments under dopamine; 19 units remained unaltered (Fig. 2A). Figure 2E shows the response of a unit to STD (blue) and DEV (red) in the control condition (left panel), during dopamine application (middle panel) and after recovery (right panel). The application of dopamine caused an increment of the STD response and a decrement of the DEV response, leading to a decrease of the CSI. In contrast, the unit in Figure 2F showed a decrement of the response to both STD and DEV during the application of dopamine, thus resulting in an increase of the CSI. The effects of dopamine peaked around 8-10 minutes after microiontophoretic application, followed by a progressive recovery to baseline values that could take beyond 90 minutes (Fig. 2E-F, right panels).

### Eticlopride effects on the CSI

We aimed to determine whether dopaminergic effects on the CSI were mediated by D2-like receptors, as suggested by previous reports [9,13]. To test endogenous dopaminergic modulation on SSA mediated by D2-like receptors, we applied eticlopride, a D2-like receptor antagonist, to 43 units. We observed no significant response changes at sample level (DEV FR: p=0.609; STD FR: p=0.769; SFR: p=0.405; CSI change: p=0.170; Figure 3A-D). However, we performed a bootstrap analysis to evaluate the statistical significance of CSI changes in each unit under eticlopride influence, which revealed that only 11 units remained unaffected. The CSI had significantly increased in 19 units and decreased in 13 units (Fig. 3A), implying that eticlopride was indeed antagonizing endogenous dopaminergic modulation mediated by D2-like receptors on those units.

### Dopamine effects on the iMM and CAS

To test whether dopamine modulates PE signaling in the nonlemniscal IC, we performed an additional set of experiments following stimulation protocol 2 (see *Methods*), which was based on the methodology of a previous study [26]. Alongside the oddball paradigm (Fig. 4A), we recorded responses of 43 units to two cascade sequences, which consisted of 10 tones presented in a predictable succession of increasing or decreasing frequencies (Fig. 4B).

**Figure 4.**
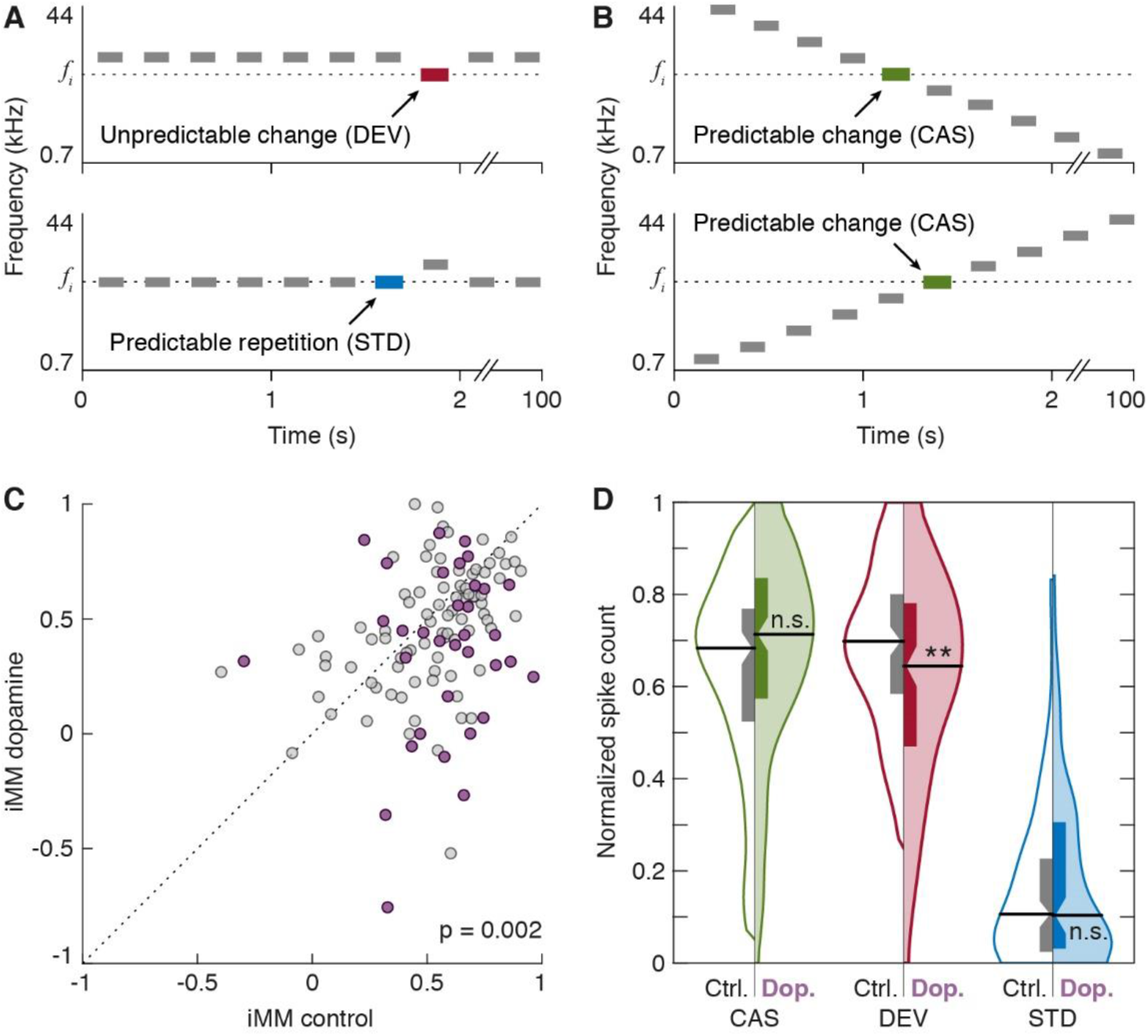
Dopamine effects on predictable versus unpredictable auditory events. **A)** Oddball paradigm, displaying two experimental conditions for a given *fi* target tone. **B)** Cascade sequences highlighting the *fi* target tone. **C)** Scatter plot of the iMM in control condition versus dopamine application. Frequencies that underwent significant iMM changes are represented in purple, whereas the rest are marked as gray dots. **D)** Violin plots of the CAS (green), DEV (red) and STD (blue) normalized responses. Control conditions are represented in the left half of each violin (no color) while dopamine effects are on display in the right half (colored). Horizontal thick black lines mark the median of each distribution, while the boxplots inside each distribution indicate the interquartile range, with the confidence interval for the median indicated by the notches. Regarding statistical significance, **n.s.** indicates that p > 0.05 and **\*\*** indicates that p < 0.01 (repeated measures ANOVA, Dunn-Sidak correction).

We used a bootstrap analysis to evaluate the statistical significance of the effect of dopamine on the iMM of each recording, which confirmed that 23 units underwent heterogeneous iMM changes (Fig. 4C, colored dots, each representing one tested frequency) whereas another 18 remained stable (Fig. 4C, gray dots). Results agreed with those obtained using the CSI, as the iMM of the sample fell by 22%, from a median of 0.57 [0.41-0.69] in the control condition to a median of 0.45 [0.27–0.65] under dopaminergic influence (p=0.002; Fig. 4C). This was caused by a significant reduction in the median DEV response (control normalized FR: 0.70 [0.58–0.80]; dopamine normalized FR: 0.64 [0.47–0.78]; p=0.002), while the STD response was not affected (control normalized FR: 0.11 [0.02–0.23]; dopamine normalized FR: 0.10 [0.03–0.31]; p=0.188). Most interestingly, CAS response also remained unaffected by dopamine application (control normalized FR: 0.68 [0.52–0.77]; dopamine normalized FR: 0.71 [0.57–0.83]; p=0.115; Fig. 5D).

## Discussion

We recorded single- and multi-unit activity in the nonlemniscal IC under an auditory oddball paradigm while performing microiontophoretic applications of dopamine and eticlopride (D2-like receptor antagonist). Following the discovery of PE signaling activity in the nonlemniscal IC [26], we included cascade sequences [44] in a subset of experiments to address dopamine role from a predictive processing standpoint. This resulted in 3 stimulation conditions: (1) STD or *expected repetition* (Fig. 4A, bottom), susceptible of generating intense SSA; (2) DEV or *unexpected change* (Fig. 4A, top), which should elicit the strongest PE signaling; and (3) CAS or *expected change* (Fig. 4B), a condition featuring the same STD-to-DEV step, but which neither undergo SSA (unlike STD) nor should entail a PE (or, at least, not as strong as DEV). Our results revealed that dopamine modulates SSA and PE signaling in the nonlemniscal IC.

### Dopamine reduces DEV responses and thereby SSA indices at population level

Dopamine application caused a 15% reduction of SSA in the nonlemniscal IC (Figs. 2A, 4C) due to general drop in DEV responses of 25% (Fig. 2B, C). Neither STD (Fig. 2B, D) nor CAS responses were significantly affected at population level (Fig. 4D). The differential effect of dopamine on DEV and STD cannot be explained by the differences in their control FR, since CAS yielded FRs as high as DEV that were not similarly reduced by dopamine (Fig. 4D). In other words, dopamine application decreased the responsiveness to *unexpected* stimuli while the responsiveness to the expected stimuli remained stable. Therefore, dopamine likely modulates net PE signaling from the nonlemniscal IC.

Eticlopride effects did not describe significant tendencies at population level (Fig. 3A-D). Nevertheless, 75% of our sample manifested significant SSA changes under eticlopride (Fig. 3A, colored dots). This confirms the release of endogenous dopamine, as well as the functional expression of D2-like receptors in the nonlemniscal IC. Taken together with previous findings regarding dopaminergic modulation of the IC [9,10,13,14], the reduction of SSA and PE signaling is most likely mediated by D2-like receptors.

The net reduction of DEV responses with dopamine is unique as compared with the effects of other neurotransmitters and neuromodulators on IC neurons. GABAergic and glutamatergic manipulations alter the general excitability of IC neurons, thereby exerting symmetrical effects on STD and DEV responses which result in a gain control of SSA [42,46,47]. Conversely, cholinergic and cannabinoid manipulation yield asymmetrical effects that mostly affect STD responses [38,39]. Activation of M1 muscarinic receptors and CB1 cannabinoid receptors tended to reduce average SSA by increasing responsiveness to repetitive stimuli (i.e., STD). Dopamine also deliver asymmetrical effects, but in contrast with the aforementioned cases, the activation of D2-like receptors tends to reduce population SSA by decreasing responsiveness to surprising stimuli (i.e., DEV).

### Intrinsic and synaptic properties generate heterogeneous dopaminergic effects

In line with previous reports [10,14], dopaminergic effects were heterogeneous across units. Complex dopaminergic interactions altering the excitation-inhibition balance cannot be accurately tracked, because the exact location and neuronal types expressing D2-like receptors in the IC are yet to be determined. Notwithstanding, the heterogeneity of dopaminergic effects must result from distinctive intrinsic and synaptic properties.

D2-like receptors are coupled to G proteins which regulate the activity of manifold voltage-gated ion channels, adjusting excitability depending on the repertoire expressed in each neuron [48]. D2-like receptors coupled to Gi/o proteins can both increase potassium currents and decrease calcium currents via βγ subunit complex, thereby reducing excitability [48]. The opening probability of calcium channels can also diminish by the activation of D2-like receptors coupled to Gq proteins [48]. D2-like receptor activation can augment or reduce sodium currents depending on the receptor subtypes expressed on the neuronal membrane [48]. Furthermore, D2-like receptor activation can also reduce NMDA synaptic transmission, decreasing the FR [48]. In addition, nonlemniscal IC neurons express hyperpolarization-activated cyclic nucleotide-gated (HCN) channels [49,50], which can be modulated by dopamine and yield mixed effects on neuronal excitability [51].

In addition, dopamine and eticlopride can interact with D2-like receptors expressed in a presynaptic neuron. Both glutamatergic and GABAergic projections converge onto single IC neurons [6,52,53], which may also receive dopaminergic inputs from the SPF [8–12]. Dopamine could potentially activate presynaptic D2-like receptors expressed in a glutamatergic neuron, as described in striatal medium spiny neurons [54–56], or conversely in a GABAergic neuron, as demonstrated in the ventral tegmental area [57].

### SPF dopaminergic projections to the IC cortices: An early precision-weighting mechanism in the auditory system?

Dopaminergic function has been traditionally studied in the context of reinforcement learning, where dopamine is thought to encode the discrepancy between expected and observed reward in a ‘reward PE’ [58,59]. Dopaminergic neurons report positive PE values by increasing their firing and negative PE values by reducing their tonic discharge rates, thereby guiding the learning process [60]. Hence, these signed ‘reward PEs’ encoded by dopaminergic neurons are substantially different from the unsigned ‘sensory PEs’ encoded by auditory neurons in the nonlemniscal IC [21,23]. According to this interpretation, our dopamine ejections in the nonlemniscal IC could mimic reinforcing signals, which in natural conditions would come from the SPF [8–12]. Speculatively, such dopaminergic input might aim to induce long-term potentiation on IC neurons to build lasting associations between acoustic cues and rewarding outcomes, thereby contributing to establish reward expectations. However, we fail to see why these positive reward PEs would mitigate the transmission of sensory PEs from the nonlemniscal IC, as evidenced by the reduced DEV responses we have observed after dopamine application.

An alternative interpretation from the predictive processing framework argues that information (i.e., PE) about the hidden states of the world cannot be encoded by dopamine release, since dopamine cannot directly excite the postsynaptic responses which would be needed to mediate the influence of that information [27]. Dopamine can only modulate the postsynaptic responses to other neurotransmitters, a function more compatible with expected precision encoding and PE weighting. This approach spares the need of two distinct types of PE signaling, while better accommodating some findings that were not easily explained as reward PEs [27]. A significant portion of dopaminergic neurons increase their firing in response to aversive stimuli and cues which predict them, contrary to how reward PEs should work [61–63]. Most relevant to the present study, dopaminergic neurons also respond to conditions where the reward PE should theoretically be zero [64,65], including novel or unexpected stimuli [66–69].

Everything considered, we propose that the most suited way of interpreting our data is through the lens of a precision-weighting mechanism. A tentative predictive processing explanation is that dopamine release in the nonlemniscal IC encodes expected imprecision. In other words, the role of dopaminergic input to the IC cortices could be to apply a net negative gain which dampens the forward propagation of PE signals. Such net inhibitory effect over sensory transmission would especially affect DEV responses, as the unexpected interruptions of an otherwise very regular train of stimuli will elicit the strongest PE signaling activity of all conditions tested in this study.

In natural conditions, such midbrain-level modulation of the bottom-up PE flow could be shaped by SPF dopaminergic neurons projecting to the IC cortices [11,13,70]. Many auditory nuclei project to the SPF, including auditory cortex, auditory thalamus, the superior olivary complex and the IC itself, providing the SPF with rich auditory information [70–74]. Even other centers, which perform higher-order functions in the sensory processing and integration of auditory information, send projections to the SPF, such as the medial prefrontal cortex and the deep layers of the superior colliculus [73]. Hence, the dopaminergic activity of the SPF must be interwoven to a great extent with the general functioning of the auditory system. The reciprocal connectivity of the SPF with many nuclei at multiple levels of the auditory pathway could provide the structural basis for an optimal encoding of expected precision, modulating via dopaminergic input the weight of PEs forwarded from the nonlemnical IC.

### Limitations

Our proposal might seem at odds with some previous works regarding neuromodulation in the predictive processing framework. Whereas cholinergic and NMDA manipulation are often reported to yield precision-weighting effects [20,30,75], dopaminergic effects on PE are less common in the literature [76], and they are usually linked to processes of active inference [27,77–80], rather than perceptual inference and learning. Besides, classic neuromodulators are often thought to increase the expected precision of PE signaling [30], contrary to the inhibitory net effects of dopamine that we found in the nonlemniscal IC.

Notwithstanding, it is important to keep in mind that the current view on the relationship between neuromodulation and expected precision derives from, and mainly refers to, cortical data. Cortical intrinsic circuitry and its neuromodulatory sources are vastly different to those of the nonlemniscal IC. Cortical predictive processing implementations have proposed specific hypotheses about the neuronal types encoding precision-weighted PEs in a defined canonical microcircuit [16,21,23]. Unfortunately, to the best of our knowledge [4,6], current understanding on the intrinsic circuitry of the nonlemniscal IC does not allow to directly import such hypotheses into auditory midbrain processing. The observed heterogeneity of dopaminergic effects in our sample is indeed compatible with distinct neuronal types fulfilling differentiated processing roles in the intrinsic circuitry of the nonlemniscal IC. However, it is not possible to distinguish between neuronal types and assign them putative roles solely based on their functional data [81]. In any case, our results make room for distinct roles of neuromodulation along the multiple stages of predictive processing along the auditory hierarchy. A possibility that could be more adequately addressed in future studies.

## Conclusions

Our study demonstrates that dopamine modulates auditory midbrain processing of unexpected input. We propose that dopamine release in the nonlemniscal IC could encode expected imprecision, consequently reducing the postsynaptic gain of PE signals and thereby dampening their drive over higher-level processing stages. The dopaminergic projections from the thalamic SPF to the IC cortices could be the biological substrate of this early precision-weight mechanism. Thus, despite being usually neglected by most corticocentric approaches, our results confirm subcortical structures as a key element of predictive processing, at least in the auditory system.

## Acknowledgments

CVB and GVC contributed equally to this work. We thank Drs Edward L Bartlett, Nell Cant, Adrian Rees and Richard Rosch for their useful comments on previous versions of the manuscript. We also thank Mr Antonio Rivas Cornejo for taking care of histological processing.

## List of abbreviatures

CAS: cascade condition.
CSI: common SSA index.
dB SPL: decibels sound pressure level.
DEV: deviant condition.
FR: firing rate (spikes/s).
FRA: frequency response area.
IC: inferior colliculus.
iMM: index of neuronal mismatch.
PE: prediction error.
SFR: spontaneous firing rate (spikes/s).
SPF: subparafascicular nucleus of the thalamus.
SSA: stimulus-specific adaptation.
STD: standard condition.
TDT: Tucker-Davis Technologies.

